# Starting from the end: what to do when restricted data is released

**DOI:** 10.1101/085100

**Authors:** Marta Teperek, Rhys Morgan, Michelle Ellefson, Danny Kingsley

## Abstract

Repository managers can never be one hundred percent sure of the security of hosted research data. Even assuming that human errors and technical faults will never happen, repositories can be subject to hacking attacks. Therefore, repositories accepting personal/sensitive data (or other forms of restricted data) should have workflows in place with defined procedures to be followed should things go wrong and restricted data is inappropriately released. In this paper we will report on our considerations and procedures when restricted data from our institution was inappropriately released.

## Data sharing at the University of Cambridge

The University of Cambridge uses DSpace as its institutional research repository which has been in place for over a decade (Smith et al. 2003). The repository was created primarily to enable open access to research publications, but has expanded in scope and now accepts a growing number of datasets (Figure 1). Thanks to intense advocacy for the benefits of data sharing, more and more researchers are prepared to share their data, including those doing research with human participants and those who need provisions for controlled access to research data. At the moment Cambridge is undertaking scoping work for the introduction of workflows and procedures to enable storing and sharing restricted data via the institutional repository. In the meantime, researchers involved in studies with human participants are advised to share resulting research data via designated specialist repositories able to handle datasets requiring controlled access.

**Figure 1.**
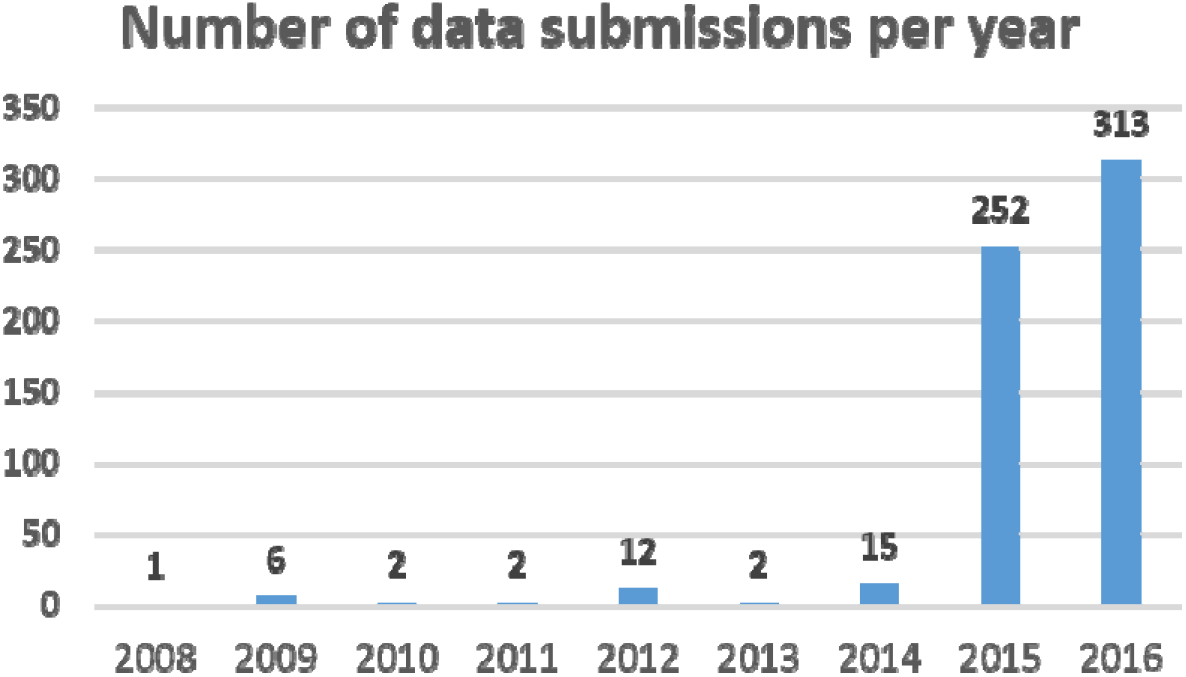
In just over a year the University of Cambridge research repository received almost ten times more data submissions than during a decade.

## Incident of inappropriate data release

Most funders now require that research data from publicly-funded research projects are submitted to data repositories within a certain timeframe from the end of the grant, regardless of whether the associated publication had already been published or not (Teperek & Kingsley 2016; Enoch 2016). During 2015, a researcher from the University of Cambridge suffered an inappropriate release of restricted research data. The dataset was held in an external data repository, specializing in storing personal/sensitive data. The released dataset was submitted to the data repository before the corresponding publication had been published and was therefore subject to a pre-publication embargo. Additionally, the released dataset was also protected by a license agreement specifying the re-use conditions.

The repository informed the researcher about the inadvertent release a couple of weeks after the repository noticed the error and after the dataset had been downloaded several times. We had to act immediately once aware of the breach to provide the researcher with appropriate support and advice on how to proceed. At the same time we were conscious that having robust workflows for dealing with this type of situations might be useful for us when setting up provisions for controlled access to data in our institutional repository. A decision to host personal/sensitive datasets in a repository entails being prepared for a situation that datasets might be inappropriately released. We undertook a risk assessment and decision-making process was thoroughly documented, which formed the basis for the workflow described below.

## Risk assessment

In order to decide on the best course of action it was essential to first analyze risks resulting from the inappropriate data release. We had to consider three different risks, which are explored below.

### Risks to study participants

The primary concern in any inappropriate data release must be to ensure that the risk of harm to the participants in the research has not increased as a result of the release. While the risks to participants will have been considered as part of the research ethics review process for the project that produced the data, approval may have been granted on the basis of particular protections or restrictions being put in place for the research data.

In this case the dataset had been anonymized, but the nature of the data had led the researcher concerned to request that access to the data be restricted. There are a number of reasons why a researcher may request that access to their data be restricted (see Box 1) (OECD 2007; UK Open Research Data Forum 2016; Corti 2011). Normally users of restricted data will be required to sign a data license agreement and may, in some cases, be vetted (for example to confirm that they are academic researchers) before the data is released to them.

#### Box 1.

##### Some possible reasons for restricting access to data

- Safeguarding participants’ data to prevent the risk of re-identification (Hobcraft 2015)
- Consent forms not allowing open sharing (Kaye et al. 2009)
- IP protection or commercial sensitivity of data (Napoli & Karaganis 2010)
- Export control laws (Relyea 2003)
- Confidential nature of research

In this case, the inappropriate release of the data had meant that users received the data without signing a required data license agreement. Data license agreements provide protection against certain potential risks to participants, for example the risk of re-identification of research participants, the inappropriate use of the data, or the release of the data beyond those for which consent for sharing was given.

We therefore approached the researcher concerned to gauge the potential risk of harm coming to participants in these ways. It became clear from these discussions that the data had been robustly anonymized and that the risk to participants of re-identification or harm through inappropriate use of the data was very low.

### Risks to the researcher

The risks to the researcher who shared the data via the repository was our next consideration. The data was embargoed as it was deposited prior to publication and changes to the dataset were anticipated (additional files were still to be added). The release of data before the end of an embargo period poses a number of potential risks to a researcher.

Firstly, embargo periods are, in part, designed to ensure that the creator of a dataset has the first opportunity to publish the results of their research. The early release of the data creates the chance that a third party could publish research on the dataset that duplicated work being undertaken by the creator of the data – potentially resulted in wasted effort on behalf of our researcher.

Secondly, we were concerned that the release of the dataset into the public domain before publication could have an impact on the researcher's ability to publish work emerging from the project. This would be particularly of concern if the dataset was published openly by an individual who had downloaded the released data.

In this case the number of downloads of the released data was low, and therefore it was judged that the risk of publications based on the data being released before the researcher published his/her results was minimal. However, we did judge there to be a risk to the researcher that publishers might raise concerns about the release of the data.

### Reputational risks

Lastly, there were reputational risks that needed to be considered, both to the researcher and the institution.

Firstly, if harm came to the participants, for example through their re-identification in the public domain, this could have a serious reputational risk for the researcher and the institution. It might also open both up to complaints or other action from the participants involved.

Secondly, if the researcher’s possibility to publish his/her work was diminished, this could make the funder of that research disinclined to fund future work by researcher (poor return on investment), therefore presenting a risk both to the researcher and to the institution.

Thirdly, the release of incomplete data posed the potential risk that re-users who undertook work on incomplete data might misinterpret them or make incorrect conclusions. This could have potential reputational risks for the researcher concerned who may be referenced as the source of the incomplete data.

The reputational risks in this case were judged to be low as the risk of re-identification was minimal and the number of downloads of the data was small. However, it was felt that actions still needed to be taken to ensure that the researcher's reputation was not affected.

## Risk mitigation

Having analyzed the risks, we then looked into strategies for risk mitigation. The best way to mitigate all the risks described would have been to contact the people who downloaded the data and ask them to sign the licence agreement. In some cases this may be possible, for example where users are required to register their contact details when downloading data. However, in this case it was not possible to contact those who had downloaded the data as the dataset had been released as fully open access files, not requiring the users to sign in with the repository. Therefore, the only information which was available to us was the IP addresses of data re-users. This information was insufficient to identify the downloaders. We therefore considered alternative strategies for risk mitigation.

### Risks to study participants

As noted above, our assessment of this incident suggested that the risk to participants was low. Following discussion with the researcher concerned, it was agreed that informing the participants about the incident would be ineffective and had the potential to cause unnecessary distress. The Research Ethics Committee who had provided the original ethical review for the project was informed of the situation.

Although no further action was taken, it was agreed that it was important that both the University Research Office and the Research Ethics Committee were informed of the situation to ensure that the inappropriate release was noted and that the situation is monitored in the future.

### Risks to the researcher

As we were unable to contact the individuals who had downloaded the data, it was not possible to inform them of the embargo and mitigate the small risk of a third party publishing results before the creator of the dataset.

The focus of the University's efforts to support the researcher, therefore, was on mitigating the risk that a publisher might not accept future papers based on the dataset. To do this, we offered the researcher the option of an official letter from the University to the publisher explaining the incident and disclosing the breach at the time of manuscript submission.

### Reputational risks

In order to mitigate reputation risks to the researcher and the institution, we initially contacted the funder of that research and informed them about the incident and the risk analysis we have performed. Reassuringly, the funder was grateful for being notified and assured us of their support for the researcher.

While it will not be possible to entirely mitigate against the potential for reputational risk from this inappropriate data release, which are entirely depended on the actions of unknown third parties, the researcher has been assured that the University will provide support to him should this be required.

## Workflow establishment

The experience of this breach has helped inform the development of effective workflows for dealing with similar incidents in the future (Figure 2). The key consideration at each step of our workflow is effective communication with relevant stakeholders.

**Figure 2.**
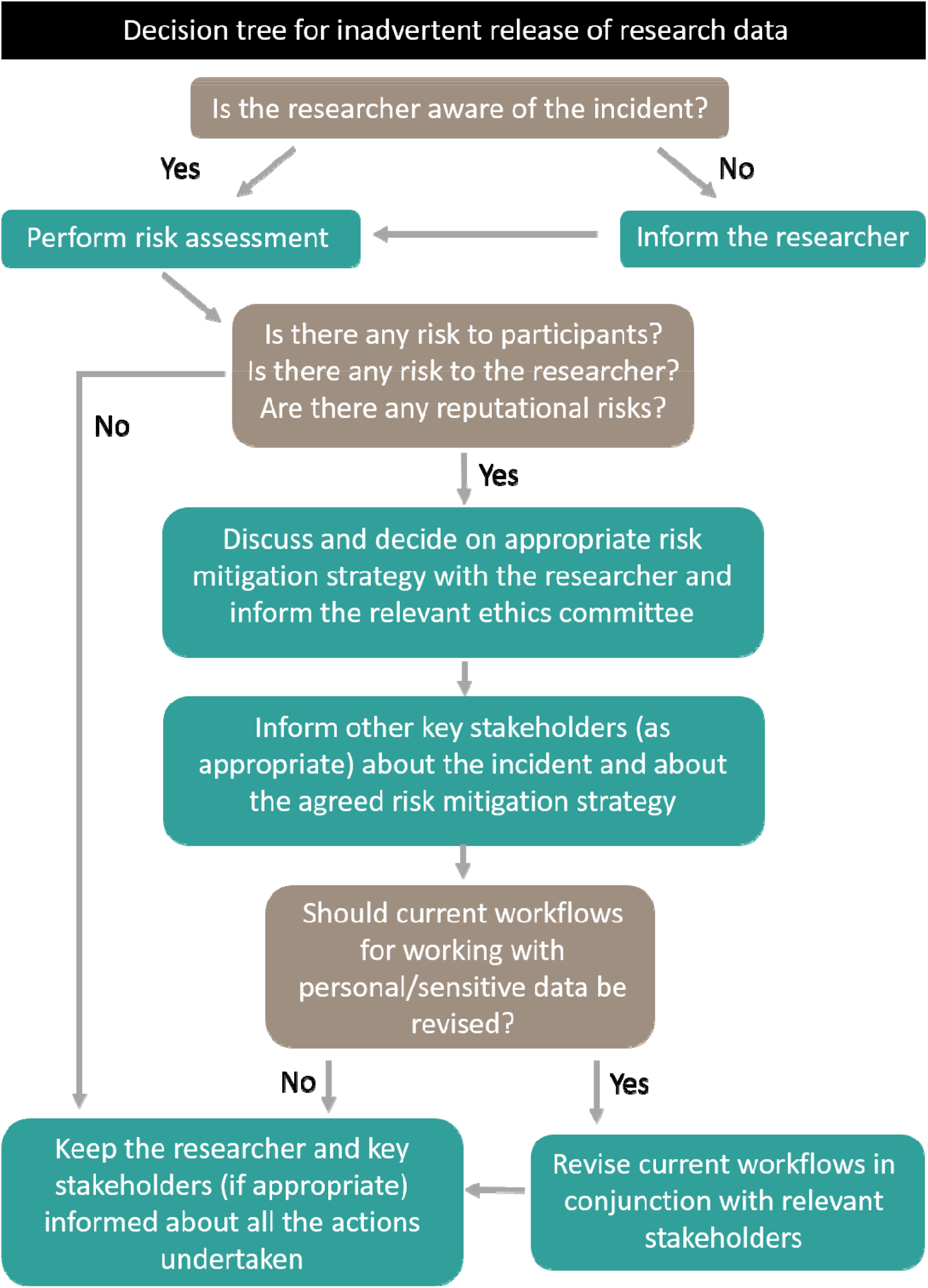
Workflow illustrating considerations when research data is inappropriate released.

## Improvements for the future

The repository team who hosted the released data reassured us that the release was due to a technical failure and the issue had been immediately solved to ensure that similar incidents do not happen in the future. However, storing and providing access to confidential data always presents a certain element of risk which needs to be assessed and managed. We decided it would be useful to analyze the situation in more detail with the view of possible workflow improvements, should similar incidents happen in the future. This was particularly important for us since we are currently developing processes for managed access to research data via our institutional repository.

### Transparency and open communication

We thought that when handling any cases when security of data was threatened, it is key to ensure transparency about the incident and open communication with all stakeholders involved. In particular, it is essential to inform the researcher whose data was released and discuss the risks involved and risk mitigation strategies as early as possible. Additionally, the following other stakeholders might need to be involved in these discussions:

- The relevant ethics committee
- The funder(s) of the research project
- Research participants
- The journal editor(s)
- People who downloaded the data
- Study administrator in the grants office

Transparency and open communication help build trust and understanding and should be treated as core values in working with personal/sensitive data.

### Guidance on working with personal/sensitive data

We also thought that the risks of participant re-identification can be substantially minimized by providing comprehensive advocacy and education program for working with personal/sensitive data. If research projects are initiated with the intention of research data being safely shared, consent forms are designed in a way to facilitate sharing and protocols for data anonymization are agreed from the start. The University of Cambridge already offers workshops on Research Integrity and Ethics, workshops on Research Data Management (Kattuman et al. 2016), as well as online guidance on creating consent forms (Morgan 2016). However, data anonymization is a more complex subject, extending far beyond simple removal of direct identifiers (Ohm 2009). Risks of re-identification are often difficult to objectively assess, especially in an era when vast amounts of information about people is available in the public domain (Ramachandran et al. 2012). Additionally, these risks evolve over time with new datasets made available and with new computational tools enabling linkage of separate datasets (Alexin 2014). Workshops on data anonymization techniques, online guidance on how to anonymize data as well as checklists for sharing anonymized data will therefore be necessary for any institution wishing to provide services for managed access to restricted data and to manage the associated risks effectively.

## Conclusions

This experience has brought sharply into focus the ever present elements of risk when depositing personal/sensitive data in a repository (even in very secure environments) and having effective workflows to mitigate them is essential for ensuring trust both among the community of researchers depositing data, as well as among participants taking part in research studies. We strongly believe that effective communication with relevant stakeholders from the earliest stages of the incident is key for optimal resolution of the situation and mitigation of possible risks.

## Acknowledgements

The authors would like to thank Dr Peter Hedges for his support and advice.

## Notes

Information about the released dataset, as well as some other information concerning the released dataset which could allow the identification of that dataset were changed.

